# Quantifying the relevance of alternative stable states in many-species food webs

**DOI:** 10.1101/2020.11.27.401711

**Authors:** Vadim A. Karatayev, Marissa L. Baskett, Egbert H. van Nes

## Abstract

Alternative stable ecosystem states are possible under the same environmental conditions in many models of 2-3 interacting species and an array of feedback loops. However, multi-species food webs might dissipate the feedbacks that create alternative stable states through species-specific traits and feedbacks. To test this potential, we develop a many-species model of consumer-resource interactions with two classes of feedbacks: specialized feedbacks where individual resources become unpalatable at high abundance, or aggregate feedbacks where overall resource abundance reduces consumer recruitment. We quantify how trophic interconnectedness and species differences in demography affect the potential for either feedback to produce alternative stable states dominated by consumers or resources. We find that alternative stable states are likely to happen in many-species food webs when aggregate feedbacks or lower species differences increase redundancy in species contributions to persistence of the consumer guild. Conversely, specialized palatability feedbacks with distinctive species roles in consumer guild persistence reduce the potential for alternative states but increase the likelihood that losing vulnerable consumers cascades into a food web collapse at low stress levels, a dynamic absent in few-species models. Altogether, among-species trait variation can limit the set of processes that create alternative stable states and impede consumer recovery from disturbance.

## Introduction

Many ecological systems exhibit distinct states characterized by the presence or collapse of consumer guilds (Estes et al. 2011). These states include pelagic food webs dominated by predators or prey (De Roos and Persson 2002; Persson et al. 2007), Caribbean tropical reefs dominated by grazing fish and corals or macroalgae (Bellwood et al. 2004; Mumby et al. 2007; Bruno et al. 2009), and salt marshes with abundant herbivores or dense vegetation (van de Koppel et al. 1996). These food web configurations might represent different states under different conditions, or they might represent the possibility of multiple alternate stable states under the same conditions, where which state occurs depends on initial conditions. Distinguishing between these two possibilities determines whether target ecosystem states recover from short-term natural or anthropogenic disturbances (Scheffer et al. 2001). Alternative stable food web states can, theoretically, arise in many ecosystems from strong feedback loops such as vegetation-dependent nutrient recycling that drives eutrophication in lakes or desertification in grasslands (Scheffer et al. 2001).

In investigations into the effect of focal feedbacks, most models predicting the potential for alternative stable states either consider 1-3 populations and omit all other taxa (De Roos and Persson 2002; Dunn et al. 2017) or aggregate species into larger guilds (Scheffer 1998; Mumby et al. 2007; May 2009). In reality, populations are embedded in larger food webs that can dissipate feedback loops across different species (Neutel et al. 2002), and heterogeneity within guilds might lead to species-specific dynamics (Lever et al. 2014). Beyond ecosystems dominated by a few strongly interacting species such as lakes, grasslands, and kelp forests (Schroder et al. 2005; Petraitis 2013), the potential for among-species differences in traits to weaken feedbacks brings the relevance of alternative stable states into question (van Leeuwen et al. 2013).

One aspect of species heterogeneity that might affect the relevance of alternative stable states is the degree to which feedbacks are species-specific (Table 1). On one extreme, aggregate feedbacks, independent of species identity, might promote multi-species alternative stable states by allowing different species involved in the feedback to help each other persist. One example of such a feedback is when the overall habitat structure provided by corals enhances herbivore grazing on macroalgae (Hoey and Bellwood 2011; Williams et al. 2001) while high aggregate macroalgal cover can impede coral recruitment (McCook et al. 2001). Coral concentration of herbivore grazing on macroalgae, with macroalgae overgrowth of corals in the absence of herbivores, can then lead to alternative coral- and macroalgal-dominated states in a two-species model (Mumby et al. 2007). At the other extreme where feedbacks affect specific, pairwise species interactions, alternative stable food web states may become less pronounced as different species collapse and recover at different disturbance levels rather than in unison. Such species-specific feedbacks can occur, for instance, in mutualistic networks where different pollinators facilitate different rather than shared plant hosts (Lever et al. 2014) and in models where strong interspecific competition is reciprocated within pairs of competitors (e.g., 7 stable states for 20 species, Levins 1966; Gilpin and Case 1976; van Nes and Scheffer 2004). Analogous feedback specialization might occur in food webs, where persistence of a consumer depends on the presence and palatability of only its primary resources. For instance, alternative stable states readily emerge in two-species models with gape-limited predation, where either (a) predators are absent and many prey survive to large body sizes or (b) predators are present and most prey are small (De Roos and Persson 2002). In multi-species systems, however, predators consuming different prey might collapse and recover at different predator mortality levels.

**Table 1:**
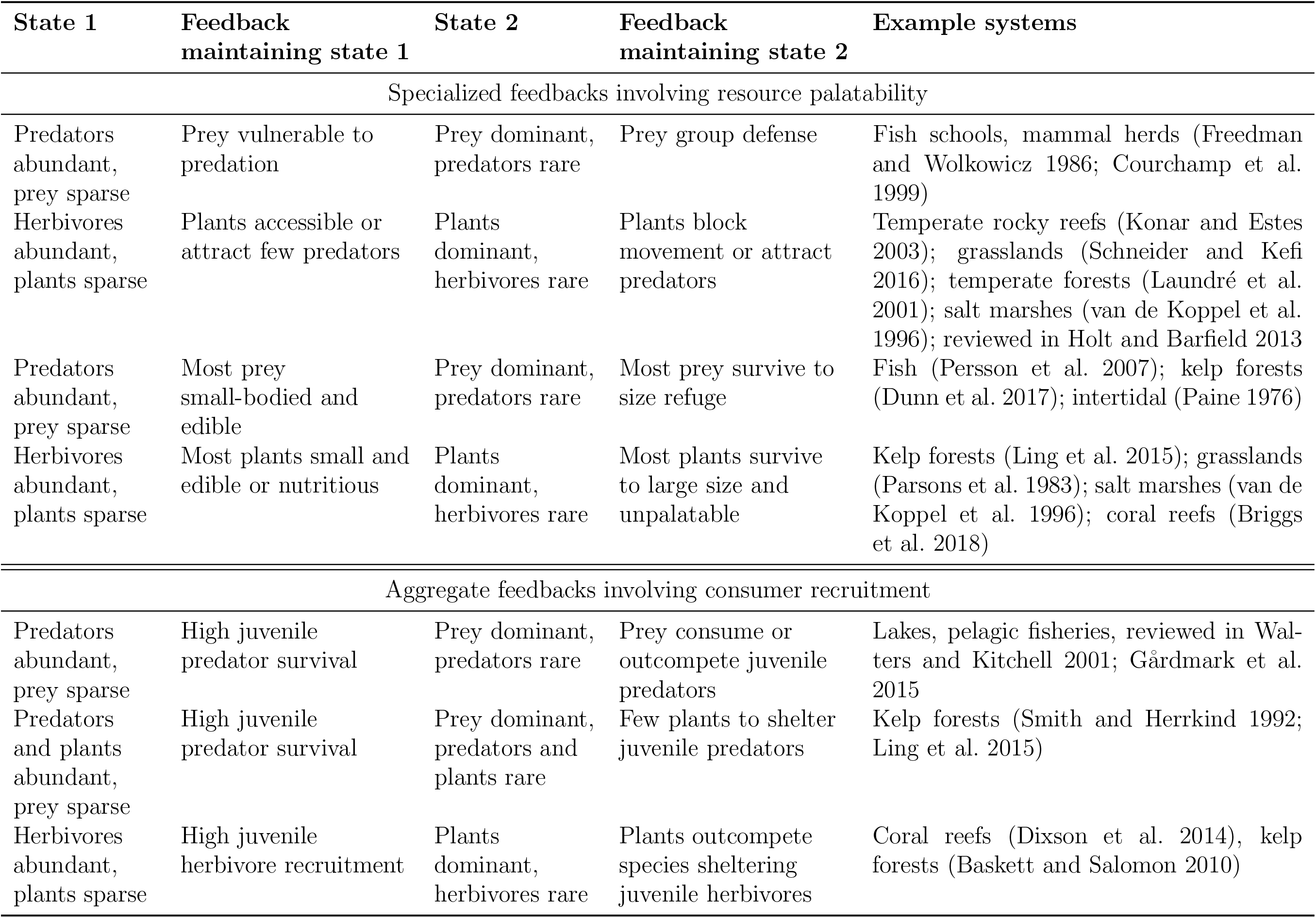
Summary of processes underlying consumer collapse via specialized feedbacks involving resource palatability (top) and aggregate feedbacks involving consumer recruitment (bottom).

The relevance of alternative stable states in multi-species food webs might additionally depend on the degree of trait diversity across species and food web connectance. While aggregated guilds and simple models might capture the dynamics of food webs with very similar species traits, trait diversity might decrease alternative stable food web states by dissipating underlying feedbacks. For instance, in the case of gape-limited predation described above, interspecific diversity in predator longevity could increase variation in harvest thresholds at which different predators collapse and recover given the greater effects of harvest in longer-lived, lower-natural-mortality species (Abesamis et al. 2014). For connectance, or the frequency of species interactions, different consumers can become increasingly interdependent as connectance increases (e.g., through a greater number of prey species per predator). Greater interdependence might either dissipate feedbacks and therefore decrease alternative states or synchronize populations to increase alternative stable states (Downing et al. 2012; Lever et al. 2014).

Here, we quantify how interactions among many species affect the potential for alternative stable food web states. For this we explore a generalized representation of two classes of models of consumer-resource interactions: aggregate or specialized feedbacks. Two-species versions of these models both exhibit alternative stable consumer- and resource-dominated states. We then examine how feedback specialization, species heterogeneity, and connectance determine the presence and distinctiveness of alternative stable states in multi-species food webs. Throughout, we focus on mortality as the driver of consumer collapse because intensive human harvest of upper trophic levels impacts ecosystems globally (Estes et al. 2011), where a key question is whether or not alternative stable states can explain delays in consumer recovery when conservation reduces harvest (Gårdmark et al. 2015).

## Methods

Our approach is to construct the simplest possible model that captures the dynamics essential to distinguishing the roles of feedback type, connectance, and trait diversity in the relevance of alternative stable states in multi-species food webs. Below we first describe the overall model structure of a food web of interacting resources and predators, where the specialized *versus* aggregated feedbacks determine the functional form of the palatability and recruitment feedbacks. We then describe how we derive those functional forms for each case of specialized and aggregated feedbacks, followed by how we implement different levels of food web connectance and trait diversity. Finally, we describe our analysis of how each factor (feedback specialization, connectance, and trait diversity) influences the prevalence of alternative stable states.

### 2.1 Food web modelling framework

Our models follow the interactions between species in a consumer guild, with densities *C_k_* for each species *k* and species in a resource guild with densities *N_i_* for each species *i*. Adult resources grow logistically at a species-specific rate *r_i_* with a carrying capacity *K_i_*. Resources experience interspecific competition with *α* as the strength of interspecific competition relative to intraspecific competition. We assume a simple hierarchy in resource competition, with greater *K_i_* values corresponding to more competitive species. Consumption follows a Type I functional response with a consumer-specific total grazing or predation rate *δ_k_*. We model diet specialization as the proportion of total consumption a consumer *k* allots to each resource *i*, Ω*_i,k_*. Each trophic interaction can additionally depend on resource palatability *P_i,k_*(*N_i_*), which represents the feedback of resource species *i* density on consumption for consumer species *k*, as detailed in the next subsection. Consumption translates into new consumer adults with a conversion constant *b_k_*. Juvenile consumer survival can additionally decline with resource abundance following 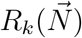, which represents the feedback of resource abundance on consumer species *k* recruitment as detailed in the next subsection. Finally, all consumers experience mortality at a density-independent rate *m* and, to help prevent competitive consumer exclusion at high connectance, morality due to intraspecific competition at a density-dependent rate *β_k_*. This produces the full multi-species model

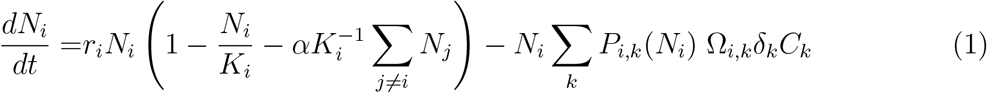

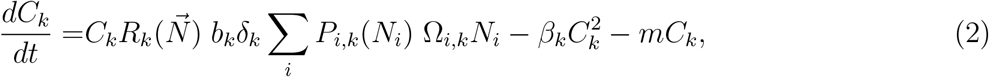

where we constrain either 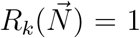 for specialized feedbacks present and aggregate feedbacks absent or *P_i,k_* (*N_i_*) = 1 for aggregate feedbacks present and specialized feedbacks absent.

### 2.2 Modelling specialized and aggregate feedbacks

We derive the predation and recruitment functions *P*_*i,k*_(*N*_*i*_) and 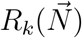 in eqns. 1–2 with either aggregate or specialized feedbacks from more mechanistic models using separation-of-time-scale assumptions. Here we summarize derivations of these simpler models, and provide mathematical derivations in Appendix A and extend our simulation results to more detailed models in Appendix B.

For specialized feedbacks, we focus on the common pattern where abundant resources are more difficult to consume (Table 1). One mechanism for this feedback, observed in both plant-herbivore (Briggs et al. 2018) and prey-predator systems (Persson et al. 2007), is size refuge from predation, with more abundant resource populations dominated by large adults that survive long enough to grow to the refuge size. A second mechanism is group defense, where grazing or attack rates decline due to prey schooling (Freedman and Wolkowicz 1986) or dense stands of plants deterring herbivory (van de Koppel et al. 1996). Therefore, the feedbacks determine how the density of resource *i N*_*i*_ affects consumption *P*_*i,k*_(*N*_*i*_) by consumer *k*.

We simplify a size-dependent predation model, wherein predation refugia and maturation occur at similar stages, to the case where juvenile resource abundance quickly reaches equilibrium on the time scale of adult resource and consumer dynamics. This assumption applies when resources have high fecundity and mature quickly. Given lower competitive ability of juveniles, juvenile abundance declines proportionally as adult abundance *N*_*i*_ approaches adult carrying capacity *K*_*i*_, causing a decline in grazing or attack rates *δ*_*k*_. For simplicity, we assume juveniles compete with or form groups with only adults of their own species in the main analysis and in Appendix C explore the case where juvenile resources additionally interact with adults of other species. Given a degree of gape limitation or group defense 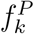, palatability of resource *N*_*i*_ to consumer *C*_*k*_ is then 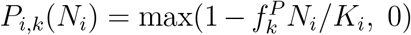 with the feedback present and 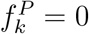, such that *P*_*i,k*_(*N*_*i*_) = 1 with feedback absent. In a two-species model, strong feedbacks impacting palatability (i.e., high 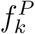) can produce alternative stable states dominated by either consumers or abundant, inedible resources over an intermediate range of consumer mortalities (Appendix A). In a food web, resources have specialized feedbacks because high density of any given resource allows that species to escape consumption, but it does not directly alter palatability of other resources (although indirect effects occur if consumer responses cascade through the food web).

For aggregate feedbacks, we focus on consumer recruitment success, which can decline with total resource abundance (Table 1). This dynamic can arise when resources reduce survival of juvenile consumers via competition or predation. In groundfish and forage fish interactions, for instance, the juveniles of different predators can be eaten their own prey (Walters and Kitchell 2001). Resources can also reduce recruitment indirectly, for instance when macroalgae overgrow corals that shelter the recruits of different herbivores (Blackwood et al. 2012). These feedbacks determine how the density of resources 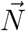. affects the recruitment 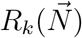 of consumer *k*.

We simplify a size-dependent consumer model, with an immature recruit stage and mature post-recruitment stage, to the case where juvenile consumer abundance quickly reaches equilibrium on the time scale of resource and adult consumer dynamics. This assumption applies when consumer fecundity is high and maturation occurs quickly. Given juvenile survival declining proportionally with resource abundance *N*_*i*_ by a factor 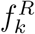, where 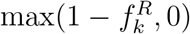 is the baseline proportion of juveniles unaffected by resources. In a food web context, resource species can differ in the degree to which they affect the recruitment of a specific consumer (see following subsection); we denote each pairwise resource *i*-consumer juvenile *k* interaction as Ψ_*i,k*_. Thus, consumer recruitment to the adult stage declines with resource density according to 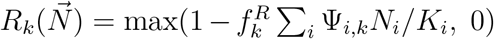 with recruitment feedbacks and 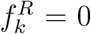, such that 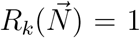 without recruitment feedbacks. In a two-species model, this feedback yields alternative stable states with adult consumers either present or extinct over a range of consumer mortality levels (Appendix A). In a food web, resources have aggregated feedbacks because they similarly impact the juveniles of multiple consumers, for instance by overgrowing a habitat-forming foundation species; for this reason, consumers collapse and resources become dominant only when aggregate resource density exceeds a threshold.

### 2.3 Manipulating connectance and trait diversity

For each influencing factor (feedback type, degree of connectance, degree of species trait diversity), we construct a series of randomly generated food webs of 12 resource and 10 consumer species. We define food web connectance as the proportion of nonzero consumer-resource interactions out of the total 120 possible interactions. For each connectance level we assign interactions randomly but omit connectance matrices where any consumer has no resources assigned. Non-zero interactions may vary in strength, for instance as abiotic conditions constrain resources and consumers to similar habitats, with Ω_*i,k*_ being the proportion of grazing rate *δ*_*k*_ a consumer *k* allots to each of its resources. To ensure a constant net strength of consumer-resource interactions across all connectance levels, we randomly draw Ω_*i,k*_ for each consumer from a Dirichlet distribution, with Σ_*i*_ Ω_*i,k*_ = 1 and *σ* representing the standard deviation of Ω_*i,k*_. As an additional, population-level metric, for each connectance level we quantify the proportion of diet comprised by each consumer’s primary resource when resources are equally abundant, averaged across all consumers (i.e., Ivlev electivity,= 10^−1^ Σ_*k*_ max_*i*_(Ω_*i,k*_)).

We account for variation in a range of species “traits” as it leads to species differences in population-level growth, competition, consumption, and life stage interactions. Among resources, we consider variation in (1) population growth *r*_*i*_ that can arise from differences in fecundity and individual growth and (2) competitive ability *K*_*i*_ that arise from differences in resource utilization efficiency. Among consumers, we consider variation in intraspecific competition *β*_*k*_, grazing/attack rates *δ*_*k*_, as well as energetic efficiency, fecundity, and juvenile survival which combine to yield *b*_*k*_. While all of these species characteristics emerge from an array of phenotypes or individual traits (e.g., body size, maturation timing, etc.), we use the term “traits” to describe them as these are how phenotypic differences affect species differences in our model.

We also model consumer species differences in sensitivity to palatability and recruitment feedbacks. Species-specific sensitivity to specialized palatability feedbacks 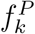 can arise from species differences in gape limitation or foraging mode, such as when prey schooling affects chase predators more than ambush predators (Turesson and Brönmark 2004). Consumer species differences in sensitivity to aggregate recruitment feedbacks can take two forms. First, consumers less affected by recruitment feedbacks overall correspond to lower 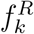 values, such as herbivore species that recruit to mangrove forests before migrating to coral reefs. Second, a resource species can dis-proportionally the juveniles of a consumer may be disproportionally impacted by a specific resource species. This example is making more sense now in terms of the actual dynamics described (the phrasing before made it sound like the reef fish were preferentially recruiting to the algae), but it still doesn’t work as an example of the point of the sentence that precedes it: the fish larvae are not competing with or being preyed upon by the algae. So, either the sentence setting up the general idea needs editing or it needs a different example that is about disproportionate competition or predation.

For example, overgrowth of exposed fore reefs by wave-tolerant macroalgal species can dis-proportionally reduce the recruitment of reef fish species that, as larvae, preferentially settle on fore reef areas (reviewed in Booth and Wellington 1998). We define *ψ* as the largest recruitment dependence of a consumer on any single resource (averaged across all consumers), where *ψ* = 1 if the juveniles of each consumer interact with only a single resource and *ψ* = 1/12 if juveniles interact equally with all resources. We then assign pairwise resource-consumer juvenile interactions Ψ_*i,k*_ from a Dirichlet distribution, where weights Σ_*i*_ Ψ_*i,k*_ = 1 and *ψ* = 0.35.

To evaluate varying degrees of trait diversity, we draw each species’ parameter values from a uniform distribution with bounds defined by *H*, the degree of trait diversity (see Table S1 for a list of parameters). Specifically, we draw each parameter j with average value 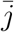 from 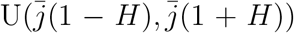. As large variations in demography did not always allow species coexistence, we omitted food webs that could not support all consumer species, i.e., where the abundance of any consumer fell below 0.1 at the lowest mortality level *m* = 0.025 given initial conditions with low resource and consumer densities. We use default or average parameter values 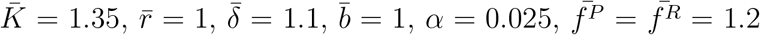, and *σ* = 0.5. We assume a low level of intraspecific consumer competition 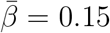 to reduce occurrence of effectively few-species food webs dominated by a few consumers, and in the Appendix we show analogous results for *β* = 0. Using this process, we generate 120 replicate food webs to analyze dynamics at each level of heterogeneity, and we generate a standard set of 120 food webs to analyze the effects of connectance.

### 2.4 Analysis of the prevalence of alternative stable states

We quantify the prevalence of alternative stable states based on two metrics: the range of consumer mortality levels producing this phenomenon and the distinctiveness of consumer- and resource-dominated states. The range of alternative stable states reflects the likelihood that initial conditions (or a short-term disturbance) affect the overall ecosystem state, whereas distinctiveness reflects whether these different states are likely to be ecologically meaningful and empirically detectable. Within this continuum, whether or not this range is zero indicates the relevance of alternative stable states.

We quantify the range of alternative stable states using a hysteresis (path dependency) analysis to find the difference between the median mortality level at which consumer species decline to extinction (abundance < 0.025) and the median mortality level at which consumers recover (abundance > 0.025). To resolve collapse points, we increase consumer mortality stepwise from 0.05 by increments of 0.02. For each step, we start at the equilibrium of the preceding mortality level and numerically simulate the model for 500 time steps to reach steady state using a Runge-Kutta method, additionally checking the final 50 time points for any cyclic dynamics. After reaching the point with all consumers extinct, we simulated improving conditions by gradually reducing consumer mortality, again with step size 0.02. To quantify recovery points, for each step we set the initial abundance of all extinct species to 0.025 to start near, but not exactly at, the equilibrium value of zero (and we start all extant species at the equilibrium of the previous mortality level). If the median mortality values for collapse and recovery are identical (range=0), then hysteresis does not occur on the community level and alternative stable food web states are not relevant. We then quantify the distinctiveness of states as the maximum difference in steady-state total consumer abundance and consumer species richness between the forward (increasing mortality) and backward (decreasing mortality) simulations across mortality values within the range of alternative stable states.

To understand how species collapses, when they occur, cascade through communities, we explore how each consumer species contributes to the persistence of the overall consumer guild. We quantify this by measuring how mean resource palatability (for specialized feed-backs) or mean consumer recruitment (for aggregate feedbacks) depend on consumer species richness. Here, more saturated relations indicate more redundant species contributions, wherein the loss of one consumer is compensated by a corresponding release from competition and increase in abundance of another consumer to maintain a consumer-dominated state. To manipulate species richness, we simulate high connectance (0.8) and high trait diversity (0.35) at low mortality (0.05), and progressively remove randomly selected consumers from the food web. After removing each consumer, we simulate the food web for 200 time steps and then measure average palatability of all resources for specialized feedbacks and average survival probability of all consumer recruits for aggregate feedbacks. We repeat this process 10 times for each food web to control for the sequence of randomly chosen extinctions, and then repeat the analysis for each of the 120 replicate food webs (i.e., 1200 total simulations for each level of species richness). One way species-specific consumer contributions to guild persistence can arise is if the extinction of one species causes secondary extinctions in a domino effect. To measure the degree to which consumers collapse in such a gradual cascade *versus* in unison, we additionally examine the transient dynamics of consumer collapse as consumer mortality exceeds the tipping point for each feedback under moderate connectance and trait diversity.

## Results

### 3.1 Effect of feedback type

Under moderate connectance and heterogeneity, alternative stable states can arise from both feedback types, but are more prevalent and distinct in the model with aggregate feedbacks (Fig. 2). Different consumers can collapse or recover at different mortality values for specialized feedbacks (Fig. 2a, 3a), but those values are simultaneous across species (with different collapse and recovery points for the full community) for aggregate feedbacks (Fig. 2b). As a result, a greater distinctiveness between states under aggregate (Fig. 2d) than specialized (Fig. 2c) feedbacks occurs throughout the range of alternative stable states. This greater distinctiveness for aggregate feedbacks holds for a range of values for connectance (Fig. 3c,d) and demographic heterogeneity (Fig. 4c,d). Greater distinctiveness with aggregate than with specialized feedbacks arises because all consumers experience the same persistence bottleneck, namely that overall resource densities remain low, in the case of aggregate feedbacks. In contrast, with specialized feedbacks each consumer can have a unique persistence bottleneck based on the edibility of its primary resource, such that each consumers can collapse at a different point depending on its unique diet.

**Figure 1:**
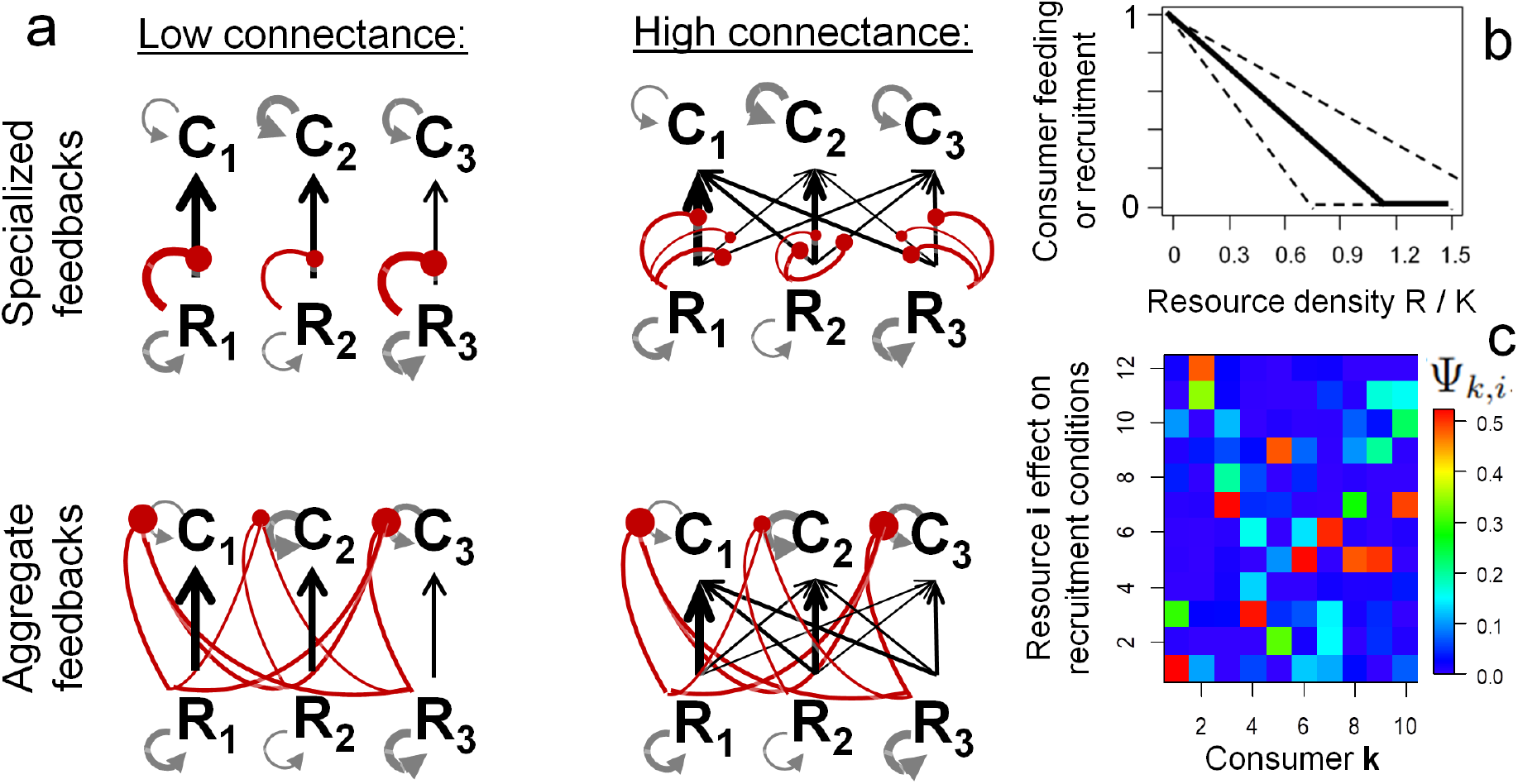
(a) Model schematic of resources (*R*_*i*_) and consumers (*C*_*k*_) showing consumption (black arrows), self-recruitment (gray arrows), and feedbacks where resources inhibit consumption in specialized feedbacks or consumer recruitment in aggregate feedbacks (red lines) for low and high levels of trophic connectance. Variable arrow thickness exemplifies trait diversity which can arise as species have different growth rates, conversion efficiency, competitive ability (thicker self-recruitment arrows), grazing rates (thicker consumption arrows), and sensitivity to resource abundance (e.g., lower gape limitation or lower reliance on corals, represented by thinner feedback lines). (b) Functional form of the feedback describing how the density of a resource, relative to carrying capacity, reduces either the resource’s vulnerability to consumption in specialized feedbacks or recruitment all consumers affected by the resource (Ψ_*i,k*_ > 0). Dashed lines indicate the maximum variation in this feedback among consumers at the highest heterogeneity levels considered. (c) Example of the matrix of weights Ψ_*i,k*_ determining the impact of resource *i* on recruitment of consumer *k*.

**Figure 2:**
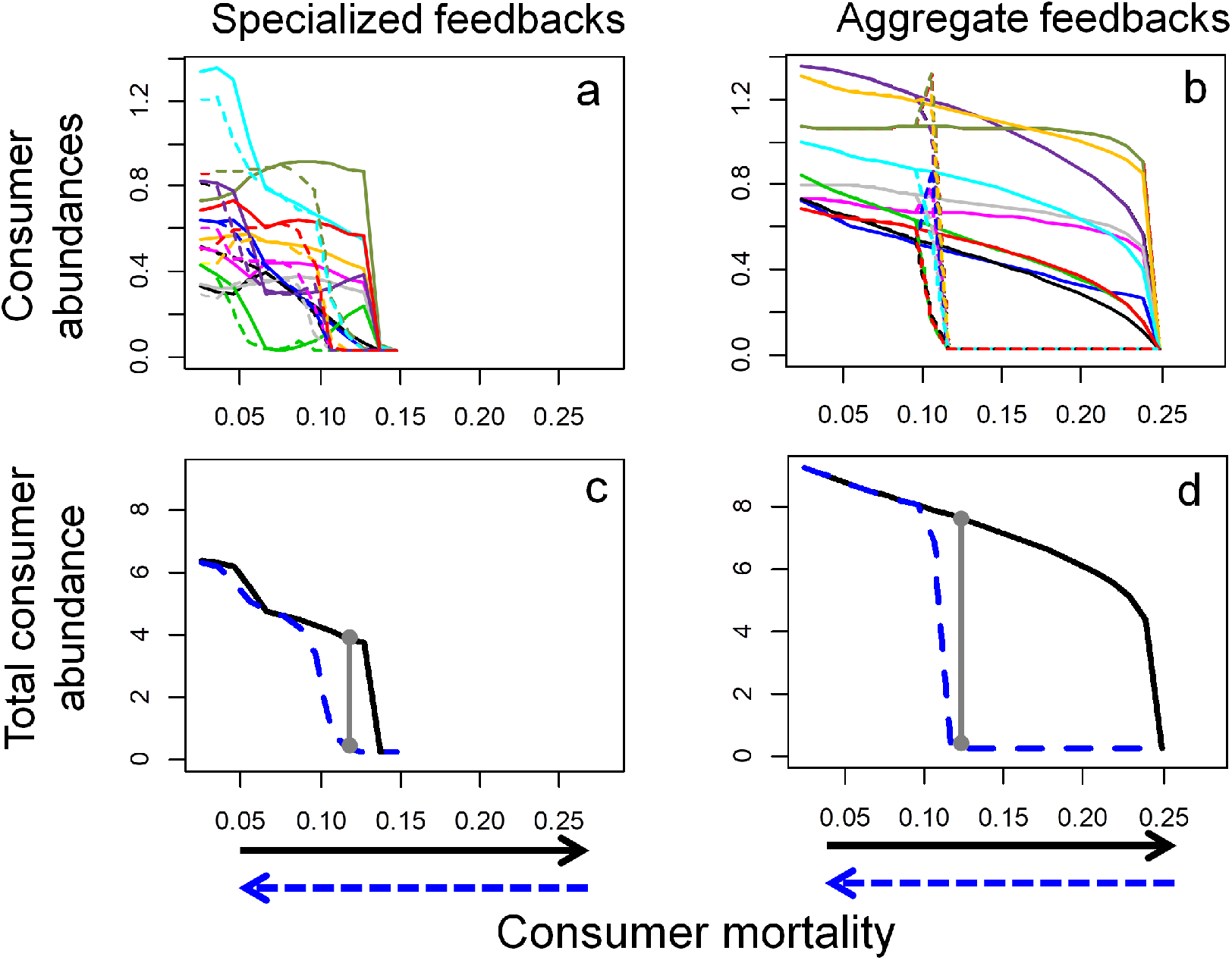
Alternative stable states are more distinct and occur over a wider range of mortality levels for aggregate (b, d) than specialized (a, c) feedbacks. Equilibrium consumer abundances as mortality sequentially increases (solid lines) and then decreases (dashed lines), shown for individual species (a, b) and total consumer abundance (c, d). Here we set connectance at 0.7 and heterogeneity at 0.15. Vertical gray lines in (c, d) show the metrics of state distinctiveness used throughout.

**Figure 3:**
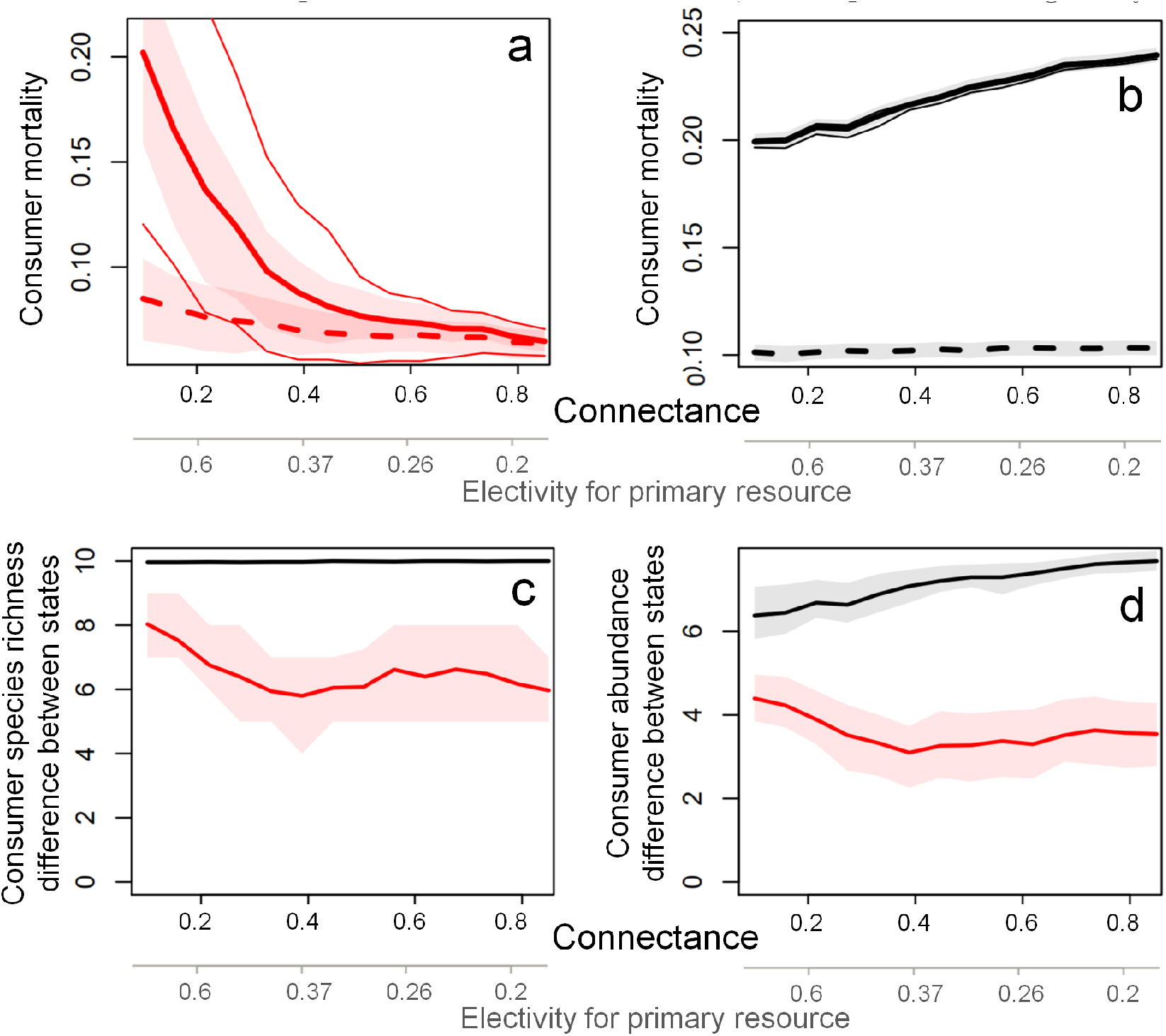
The potential for alternative stable states decreases with connectance for specialized feedbacks but increases with connectance for aggregate feedbacks. (a, b) Median thresholds at which consumer species collapse (solid lines) and recover (dashed lines) for specialized feedbacks (a, red colors) and aggregate feedbacks (b, black colors), with alternative stable states present between these thresholds. Shaded regions in (a, b) denote the interquartile range of each threshold, and thin lines denote thresholds at which the first and last consumers collapse as mortality gradually increases (variation not visible for aggregate feedbacks). (c, d) Distinctiveness of consumer- and resource-dominated states, measured in species richness (c) and in total consumer abundance (d), with shaded regions denoting interquartile ranges. The secondary x-axes denote the proportion of diet comprised by each consumer’s primary resource when resources are equally abundant (Ivlev electivity). Results at each connectance level span 120 simulated food webs, with species heterogeneity at 0.15.

**Figure 4:**
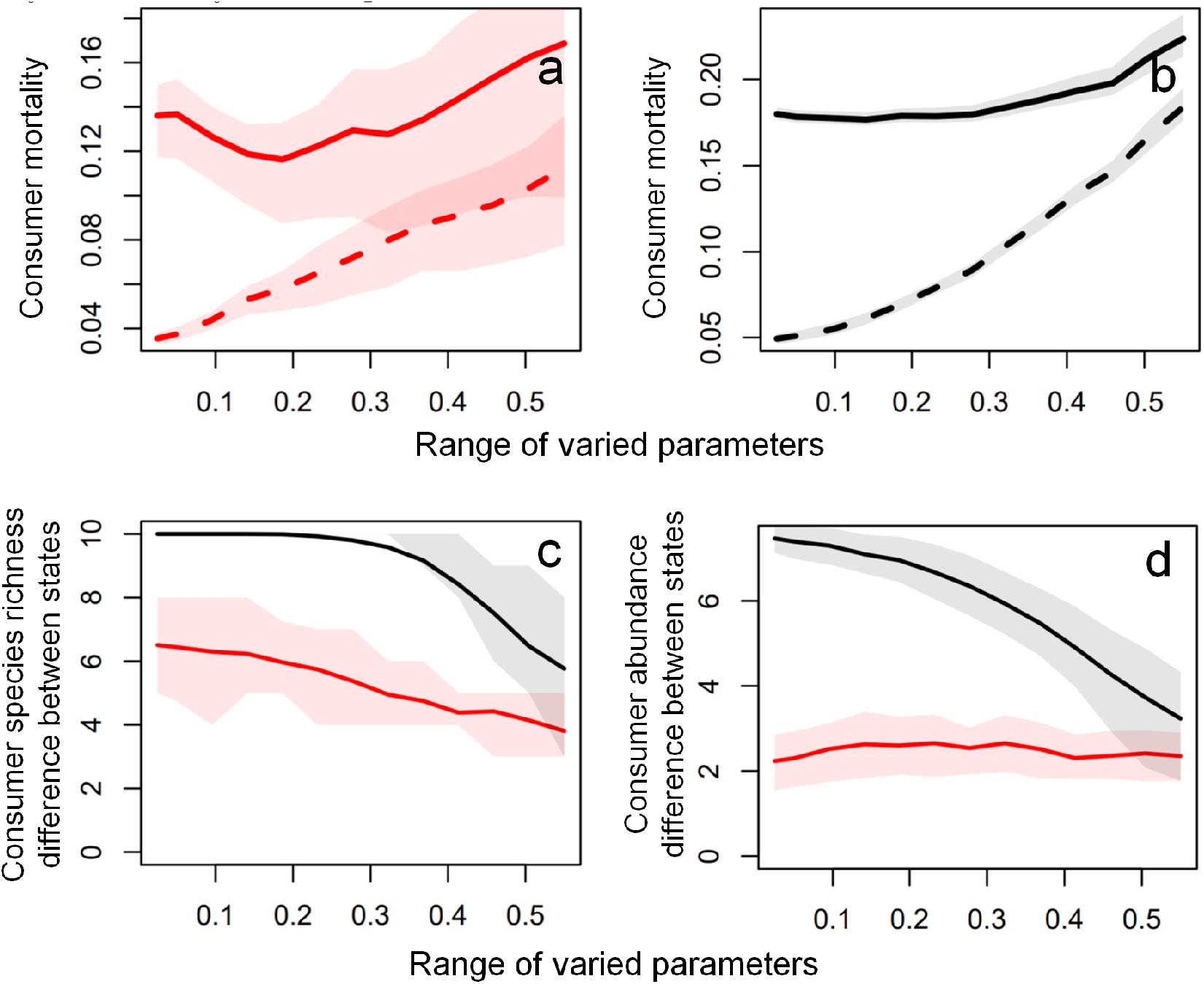
Demographic heterogeneity decreases hysteresis and the distinctiveness of alternative stable states. (a, b) Median thresholds at which consumer species collapse (solid lines) and recover (dashed lines) for specialized feedbacks (a, red colors) and aggregate feedbacks (b, black colors), with alternative stable states present between these thresholds. (c, d) Distinctiveness of consumer- and resource-dominated states, measured in species richness (c) and in total consumer abundance (d). In all panels, shaded areas denote interquartile ranges across 120 simulated food webs at each connectance level, with connectance at 0.25. Interquartile ranges of thresholds not visible in (b) because collapses and recoveries each happen synchronously across species.

Alternative stable states can also occur over a wider range of mortality values under aggregate (Fig. 2d) than under specialized (Fig. 2c) feedbacks. At low connectance, lower prevalence of alternative stable states occurs with specialized feedbacks because consumer-resource pairs exhibit separate dynamics. At moderate-to-high connectance, the escape of one resource from consumption reduces resource uptake for all species that consume the resource. This causes a decline in abundance for multiple consumers, allowing the next resource to escape consumption and repeating this feedback. In other words, the entire consumer guild can experience multiple persistence bottlenecks, where most resources must be edible for consumers to persist. We therefore find that above low connectance, the collapse threshold of the entire consumer guild converges on the threshold of the most vulnerable consumers, namely, the species that collapse at the lowest mortality levels (Fig. 3a). This arises because consumers divide their consumption among many different resources and cannot singlehandedly maintain all consumed resources in edible, low-density states. Loss of vulnerable consumers at low mortality levels and escape of their primary resources to high abundance therefore translates to reduced resource uptake for other consumers and, potentially, a multi-species, cascading series of consumer collapses (Fig. 5b,c). Accordingly, average resource palatability and the strength of specialized feedbacks maintaining the consumer guild both decline proportionally with the number of consumer species present (Fig. 5a). Aggregate feedbacks produce alternative stable states over a larger mortality range because favorable recruitment conditions require control of only the overall resource abundance. This redundancy in consumer contributions to recruitment is reflected in a weaker effect of species loss on aggregate compared to specialized feedbacks at high diversity (Fig. 5a) and causes a more synchronized loss of all consumers at the collapse point (Fig. 5b). In summary, alternative stable states are more prevalent under aggregate than specialized feedbacks because aggregate feedbacks both synchronize consumer collapse and remain strong when vulnerable species decline.

**Figure 5:**
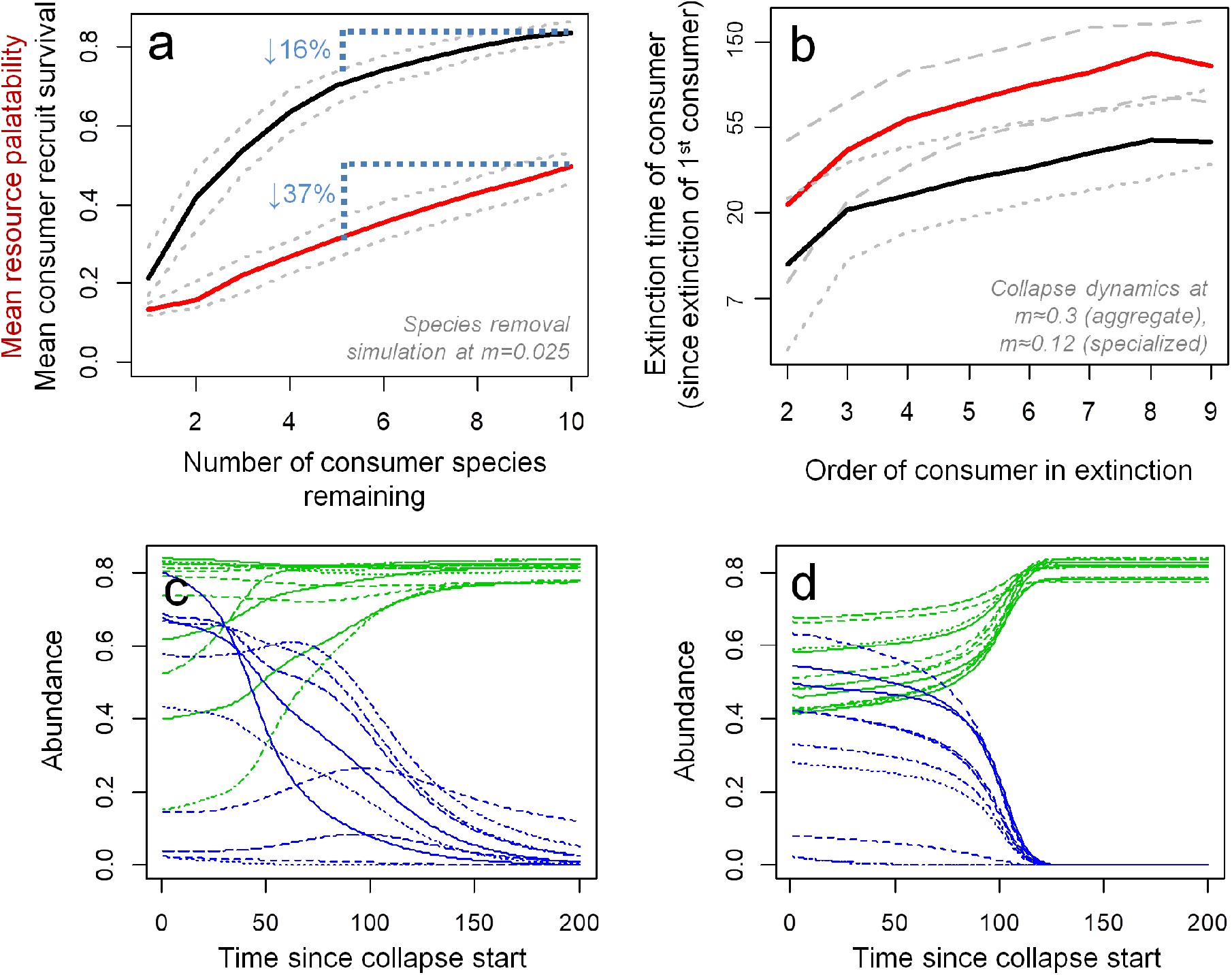
Compared with aggregate feedbacks, specialized feedbacks lead to species-specific consumer contributions to guild persistence and prolonged cascades of secondary extinctions following the loss of vulnerable consumers. (a) Effects of progressively removing randomly selected consumer species at low mortality (*m*=0.025) on the average palatability of resources (specialized feedbacks, red line) and the average survival of consumer recruits (aggregate feedbacks, black line), with increasing numbers of species removed from right to left along the x-axis; blue dotted lines denote how removing 5 consumers reduces palatability much more than recruitment. (b) Distribution of consumer extinction times, measured in time steps from extinction of the first consumer, in simulations where mortality crosses the threshold leading to the collapse of all remaining consumers, with greater values reflecting a greater frequency of cascading, delayed extinctions. (c,d) Examples of dynamics summarized in (b) showing the time series of consumer collapse (blue lines) and resource growth (green lines) over the course of consumer extinction for specialized (c) and aggregate (d) feedbacks. Lines in (a,b) denote medians (red and black) and interquartile ranges (gray lines) across 1200 simulations; note the log-scale y-axis in (b). Resource abundances in (c,d) scaled relative to carrying capacity of each resource. In (b-d), heterogeneity is 0.15 and connectance is 0.7.

### 3.2 The effect of connectance

Connectance can increase the potential for alternative stable states with aggregate feedbacks but decrease the potential for alternative stable states with specialized feedbacks (Fig. 3). For specialized feedbacks, connectance decreases the range of mortality levels leading to alternative stable states by making persistence of the consumer guild increasingly reliant on the presence of the most vulnerable species (as described above). For aggregate feedbacks, connectance increases both the distinctiveness and the range of alternative stable states across consumer mortality levels (Fig. 3b, c). This greater range with connectance arises because more dominant consumers (e.g., those with higher grazing rates or conversion efficiency) consume a larger fraction of resources, thereby strengthening the feedback loop maintaining consumer-dominated states (Fig. 3d). Greater feedback strength also allows consumers to persist at higher mortality levels but has no effect on consumer recovery from the resource-dominated state with poor consumer recruitment conditions, leading to an overall increase in the range of alternative stable states (Fig. 3b).

### 3.3 The effect of demographic heterogeneity

Given moderate connectance (0.25), heterogeneity in species demography reduces the distinctiveness and range of alternative stable states, with greater sensitivity to heterogeneity under specialized than under aggregate feedbacks (Fig. 4). Greater heterogeneity translates to the most vulnerable consumers being more vulnerable, which, following our preceding results, disproportionally weakens specialized feedbacks. In addition, alternative stable states driven by both feedbacks decline at high heterogeneity, where more dominant consumer species (e.g., those with greater consumption rate *δ*_*k*_ and conversion efficiency *b*_*k*_) can singlehandedly maintain favorable resource edibility or recruitment conditions. Such species also help other consumers recover, increasing the median recovery threshold.

## Discussion

We have shown that alternative stable states can occur in interconnected food webs of many different species, especially with aggregate feedbacks (Fig. 2) which can arise from a wide range of ecological processes affecting consumer recruitment (Table 1). A greater potential for alternative stable states through aggregate feedbacks parallels work modeling more nested mutualistic networks where multiple pollinators additively benefit a shared host (Lever et al. 2014) and community resistance to invasion by a shared competitor (Case 1990) or predator (Downing et al. 2012). In all cases, multi-species alternative stable states arise despite species differences because species help each other persist by improving conditions in the same way, for instance herbivores promoting habitat-forming corals by limiting total macroalgal cover. We find that such aggregate feedbacks translate to a greater redundancy in species contributions to maintaining favorable recruitment conditions (Fig. 5a), a result analogous to saturating relationships between biodiversity and ecosystem functioning (Schwartz et al. 2000). This functional redundancy makes the overall consumer guild resistant to the decline of sensitive consumer species that comes with increased mortality, but it does not affect recovery after the entire guild collapses (Fig. 3b). This leads to a potential resistance-resilience tradeoff (Downing et al. 2012): a community with greater functional redundancy and resistance to stress (here, collapsing at higher mortality) may also exhibit a greater potential for alternative stable states where, when it collapses, the community requires a larger reduction in stress in order to recover (i.e., lower ecological resilience *sensu* Holling 1973).

In contrast to the case with aggregate feedbacks, we find that the potential for alternative stable states declines with specialized palatability feedbacks where a resource species can become inedible to multiple consumers at high abundance (Fig. 2; Table 1). In comparison to aggregate recruitment feedbacks, where resource species can only reach high abundance in unison once most consumers decline, under specialized feedbacks each consumer species uniquely benefits the larger guild (Fig. 5a). For example, each consumer could prevent its primary resource from surviving to large body sizes and becoming inedible to other species (De Roos and Persson 2002; de Roos and Persson 2013). Loss of vulnerable species at low mortality levels therefore can reduce ecosystem functions maintaining the consumer guild (here, resource palatability) and cascade into a many-species collapse (Fig. 3a, 5). Thus, with palatability feedbacks, food web connectivity can impart the vulnerability of a few species onto the rest of the food web. In few-species models, palatability and recruitment feedbacks exhibit a similar potential to drive alternative stable states (Gårdmark et al. 2015; van de Leemput et al. 2016, Appendix A). Our multi-species simulations therefore highlight how species heterogeneity narrows but does not eliminate the set of processes that could create alternative stable states and impede food web recovery from disturbance.

Despite reducing the potential for alternative stable states, we find that specialized feedbacks can still drive multi-species collapses following small increases in mortality (Fig. 2, 5c). This result parallels spatial regime shifts where heterogeneity favors a more resilient (here, resource-dominated) state, which propagates across space from more to less vulnerable patches in a domino effect (van Nes and Scheffer 2005). In van de Leemput et al. 2016, alternative stable states present in well-mixed systems are greatly reduced when matter or organisms move slowly and affect feedbacks only in nearby locations. Localized spatial feedbacks parallel specialized feedbacks in our model, where palatability of a resource depends on intraspecific interactions (competition or group defense) and affects only the resource’s consumers, such that consumer collapse cascades gradually throughout the food web. This suggests an opportunity for future research, using models developed here as a starting point, to expand food web resilience theory based on insights from the better-developed literature on spatial regime shifts (van Nes and Scheffer 2005; van de Leemput et al. 2016) and spatial early-warning signals (Dakos et al. 2011).

### 4.1 Drivers of feedback specialization

The reduced potential for alternative stable states and reduced resistance of species guilds to mortality found here for specialized feedbacks can apply to a wide range of feedbacks and systems. These include trophic interactions in coastal and pelagic marine systems, lakes, and forests (Table 1). We point out that our results are also robust to more complex food web interactions (Appendix C), such as when the juveniles of one resource face competition from adults of multiple resource species (i.e., intraguild predation among prey or priority effects among plants) and when prey form multi-species herds or schools (Gil et al. 2018). Beyond food webs, co-occurring species can also facilitate each other in competition-dominated systems: animal species richness promotes habitat-forming foundation species when animals experience strong intraspecific competition (invertebrates cleaning sediments off corals, Stier et al. 2012; fish protecting anemones, Schmitt and Holbrook 2003), or when animal species prevent different taxa from fouling a foundation species (Stachowicz and Whitlatch 2005). In mutualistic systems, multiple pollinators often benefit a shared host plant (Lever et al. 2014). Reviewing existing case studies, Afkhami et al. (2014) found that the presence of multiple mutualist partners in most cases synergistically promoted the central host species, indicating that the mechanisms by which species indirectly facilitate each other in many systems may be species-specific.

### 4.2 Model assumptions and results robustness

For tractability and to focus on the role of different feedback types, we omitted several factors that might affect the potential for alternative stable states in multi-species food webs. Our separation-of-time-scale assumptions of high fecundity and fast maturation do not qualitatively alter our findings (Appendix B); however, explicit stage structure may reduce the potential for alternative states compared to stage-implicit models by de-stabilizing consumer-dominated states. Our approach might also overestimate this potential by assuming that feedbacks strongly affect most species. In reality, species less affected by feedback processes might reduce the potential for alternative stable states: for instance, larger predators that more readily consume large prey might prevent a prey-dominated state in which more gape-limited predators are extinct (van Leeuwen et al. 2013). Alternatively, our randomized parameter selection may underestimate the presence of alternative stable states by omitting trade-offs in species life histories. For example, on tropical reefs larger-bodied herbivores with less gape limitation (Briggs et al. 2018) or less sensitive juvenile stages might also have lower fecundity, here modeled implicitly as a lower conversion efficiency. For such species, a more limited effect of low coral cover might trade off with greater sensitivity to mortality. The capacity for heterogeneity to reduce the potential for alternative stable states emphasized here underscores the importance of future system-specific studies that model observed species heterogeneity and its underlying life history trade-offs. Our models also omit spatial structure, more complicated functional responses (e.g., Type II or III), and stochasticity, all of which could further obscure alternative stable states or reduce the range of conditions over which this phenomenon occurs (e.g., Guttal and Jayaprakash 2007). Finally, while we base our analyses on 120 randomly constructed food webs, the effects of feedback specialization, connectance, and demographic heterogeneity in any particular food web may differ from the overall patterns.

Additional factors could also increase the potential for alternative stable states with palatability feedbacks from our main analysis. In particular, consumers might adjust their diet composition to concentrate feeding only on resources which remain palatable. Further modeling work is needed to determine whether alternative stable states become more likely as this process strengthens consumer-resource interactions or less likely as converging diets increase competitive exclusion among consumers. However, we note that diet flexibility may be limited by seasonality, spatial structure, or nutrient requirements by the consumer.

### 4.3 Anticipating alternative stable states in food webs

Our results suggest that the potential for simple models to predict alternative stable states in diverse food webs depends on the relative strength of aggregate *versus* specialized feedbacks. Establishing the presence and conditions leading to alternative stable states based on field data alone is rarely feasible due to the need for long-term data of each state at large spatial scales under identical environmental conditions (Petraitis and Dudgeon 2004; Petraitis 2013). Simple data-driven mechanistic models therefore represent a common approach to quantifying the potential for alternative stable states to explain observed patterns in diverse ecosystems (Ives et al. 2008; Mumby et al. 2007, 2013; Karatayev et al. 2021). Our results show that the results of such simple models might be robust to food web complexity with aggregate feedbacks. In the presence of specialized feedbacks and trait diversity, however, accounting for additional diversity alter expectations for anticipating community collapse and alternative stable states.

In reality, many food webs have both specialized and aggregated feedbacks. On coral reefs, for example, abundant macroalgae can simultaneously (i) become less edible in dense plant stands (Briggs et al. 2018) and (ii) overgrow corals that facilitate recruitment of many herbivores (Dixson et al. 2014). This raises two avenues for future study. First, more empirical studies might quantify the strength of specialized *versus* aggregate feedbacks, with alternative stable states being more likely in systems where aggregate feedbacks predominate. Second, future theory might resolve the dynamics of systems where specialized and aggregate feedbacks co-occur.

## Acknowledgments

We thank Alan Hastings, Sebastian Schreiber, Marten Scheffer, and Mohammad Shojaei for insightful discussions that improved the manuscript. VAK was supported by an NSF Graduate Research Fellowship.

## Data and code availability

Code to reproduce all simulations available at github.com/VadimKar/Food-web-resilience-model

## Appendix A Model derivations

## 1.1 Specialized feedback model

To derive the consumer-resource model of specialized feedbacks (eqns. 1–2; see Table S1 for all parameter definitions), we start with a general model tracking abundance of consumers *C* and a resource with juvenile *J* and adult *A* stages. Throughout, we assume juveniles as being competitively inferior to adults, for instance seedlings and small plant stages that only grow on free space. Thus, adults *A* produce offspring at rate *r*, and a proportion 1 – *A* of offspring successfully recruit to the juvenile stage; note that adults may cause juvenile mortality when adults exceed carrying capacity (e.g., trees shading out seedlings). Juveniles then mature at a rate *γ*(*A*) that can decline with adult abundance (e.g., if higher adult abundance reduces resource availability for juvenile growth). Additionally, both juveniles and adults loss to consumption at a rate *δ*. Resource loss to consumption may differ between juvenile and adult resources by a factor *g*, with *g* < 1 indicating lower consumption of adults, as might arise from consumer gape limitation. Finally, consumers have a conversion efficiency *b* and mortality rate *m*, giving the full dynamics:

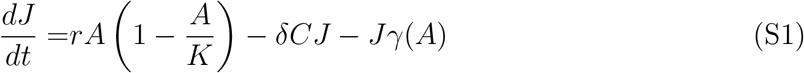

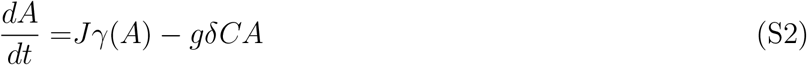

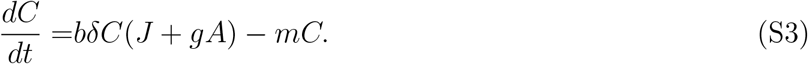

We simplify these dynamics first by assuming that juvenile resource stages are relatively brief on the time scale of adults and consumers, that is, *r, γ*(*A*) *>> δ*, which yeilds

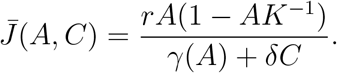

Second, we simplify the consumption term (*γ*(*A*)+*δC*)^−1^ using the first two terms of a Taylor approximation expanded with respect to *C* around *C* = 0 to arrive at a consumer-resource interaction form where juvenile abundance declines linearly rather than geometrically with consumer abundance. Juvenile abundance is then approximately

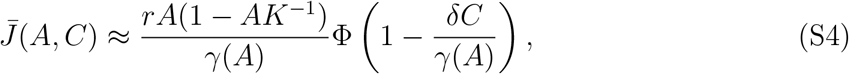

where as in the main text, we use Φ(*x*) = max(*x,* 0) to ensure that total consumption is non-negative. While differing qualitatively at high consumer densities, this simplification can approximate dynamics involving collapse of consumers from low-moderate abundance levels. Substituting this term for *J* in adult and consumer dynamics yields

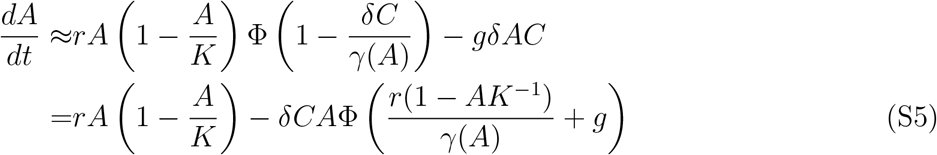

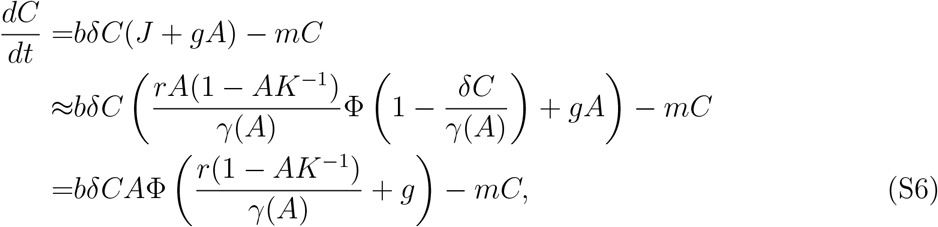

where the final simplification reflects the time scale separation *γ*(*A*) ≫ *δ*, such that *δ/γ*(*A*) ≈ 0.

In the case of density-independent maturation rates (i.e., *γ*(*A*) = *γ*_0_), renaming *N =A* and defining *δ′ = δ*(*g + r/γ*_0_) and *f*^*P*^ = *r*/(*r + gγ*_0_) ≤ 1, simplifies eqns. S5–S6 into eqns. 1–2:

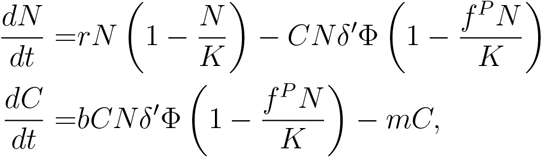

where the proportion of edible resources Φ(1 – *f*^*P*^ *N/K*), hereafter defined as *ξ*(*N*), declines with abundance. Here, note that greater palatability of adults *g* translates to a lesser decline in *ξ*(*N*) with *N* (i.e., lower *f*^*P*^), weakening the feedback that drives alternative stable states.

To explore the case where juveniles mature slower when adults deplete resources, we consider resource palatability in eqn. S5–S6 for a simple case where *g* = 0 and *γ*(*A*) = *γ*_0_(1 – *zA*), where the proportion *z* < 1 reflects maturing juveniles being less vulnerable to competition from adults than newly recruiting juveniles. In this case resource palatability is convex and predominantly declines at high adult abundance (Fig. S1a). This reflects the case where slower maturation due to resource depletion by adults translates to a greater period of resource individuals being vulnerable to predation. Analogous patterns arise at steady state in more detailed, biomass-based models (e.g., de Roos and Persson 2013, Chapter 3, eqn. 24) that explicitly model juvenile dynamics, energy uptake, and competition among adults and juveniles (Fig. S1b). Here, the proportion of total resource biomass comprised by smaller juveniles more vulnerable to consumption similarly declines in populations that experience lower juvenile mortality and reach greater total biomass. Finally, we note that in conventional models of group defense (‘Type IV’ functional forms, Freedman and Wolkowicz 1986; Bate and Hilker 2014) resource palatability can decline in a concave fashion (Fig. S1a).

## 1.2 Aggregate feedback model

Here, we derive the general model of aggregate feedbacks (eqns. 1–2) from models where resources inhibit consumer recruitment directly or indirectly.

## 1.2.1 Cultivation effects

For cultivation effects where prey *C* directly outcompete or consume juvenile predators *J* and are consumed by adult predators *P*, we start with the model of Baskett et al. (2006) (eqns. A1-A3). This model includes competitive effects of prey on juveniles *α*_*CJ*_, juveniles on prey *α*_*JC*_, and competition among prey *α*_*CC*_ and among juveniles *α*_*JJ*_. Juvenile abundance grows with predator consumption of prey (with a rate *δ*_*C*_ and conversion efficiency *β*_*C*_) and of juveniles (with a rate *δ*_*J*_ and conversion efficiency *β*_*C*_), juveniles mature at a rate *γ*, and adults die at a rate *m*. Modifying the model to have a shared carrying capacity term *K* (*K*_*J*_ = *K/α*_*JJ*_, *K*_*C*_ = *K/α*_*CC*_) gives

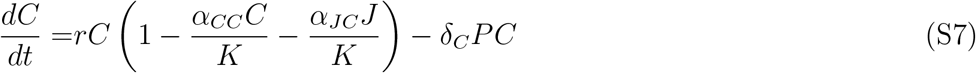

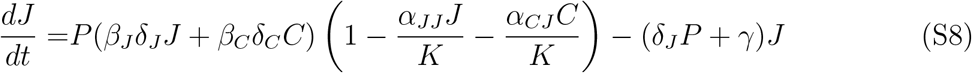

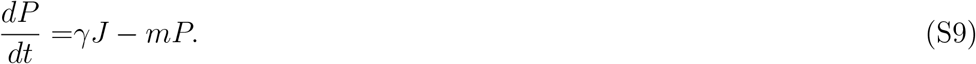

We first simplify the model by assuming negligible predator cannibalism (*δ*_*J*_ = 0), that juveniles have no competitive effect on prey (*α*_*JC*_ = 0), and negligible competition among juveniles (*α*_*JJ*_ = 0). This assumption reflects the case where juvenile body size (and consequently resource consumption) is smaller than that of prey, and may be biologically inaccurate in pristine systems with very high juvenile predator abundance. Second, we assume that juveniles can quickly mature to become adults (*γ* >> 0) and a high adult reproductive potential (*β*_*i*_ >> 0) such that juvenile abundance reaches steady state faster than prey or predator abundance. With these simplifications, steady state juvenile abundance is

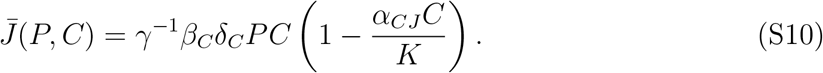

Substituting this expression into predator dynamics and setting *K′* = *K/α*_*CC*_, *δ* = *δ*_*C*_, *b* = *β*_*C*_, and *f*^*R*^ = *α*_*CJ*_ *α*_*CC*_ yields the final set of equations

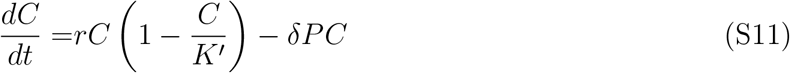

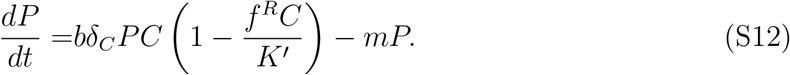

These dynamics are equivalent to eqns. 1–2 in our main analysis, where larger *f*^*R*^ corresponds to stronger competitive effect of prey on juveniles.

## 1.2.2 Consumer habitat loss effects

In addition to directly inhibiting consumer recruitment, resource species can inhibit recruitment indirectly by negatively affecting species on which consumers rely. We examine one such case where herbivores consume plants that promote survival of juvenile predators (Karatayev and Baskett 2020). Such dynamics occur, for instance, on rocky temperate reefs in California where urchins *U* consume kelp *A*, while kelp provide shelter to juvenile predators *P*, sheephead and spiny lobsters. Intense fishing of predators frequently leads to high urchin densities that overgraze kelp, causing predator collapse and producing persistent urchin barrens with few predators. Predator recruitment therefore depends on the amount of urchins consumed and increases proportionally with kelp abundance by a factor *f*_*c*_, where 1 – *f*_*c*_ is the baseline predator recruitment success without kelp. Accounting for kelp density dependence, Type I grazing rates on algae *δ*_*A*_ and urchins *δ*_*U*_, conversion constants *a*, *b*, and predator mortality *m* yields:

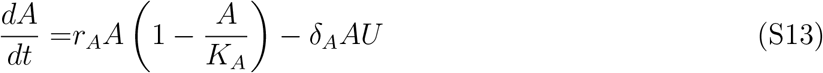

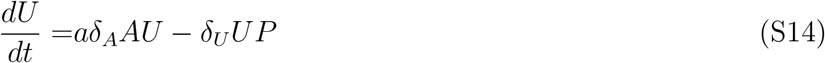

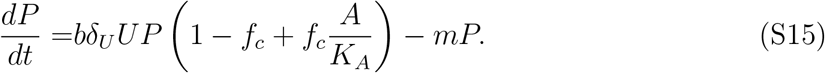

We simplify these dynamics to reflect the fact that urchins can overgraze kelp forests within weeks (*δ*_*A*_ >> 0), while at low urchin densities kelp grow rapidly and reach carrying capacity within several months (*r*_*A*_ >> 0). As a result, kelp rapidly reach steady state abundance *Ā*(*U*) = *K*_*A*_(1 – *Uδ*_*A*_/*r*_*A*_) on the time scales of urchins and predators. Plugging this term into urchin and predator dynamics yields

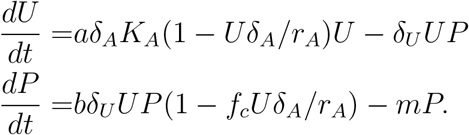

Substituting *K* = *r*_*A*_/*δ*_*A*_, *r* = *aδ*_*A*_*K*_*A*_, *f*^*R*^ = *f*_*c*_, and *δ* = *δ*_*U*_ gives eqns. 1–2:

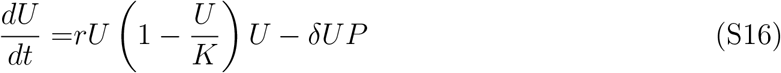

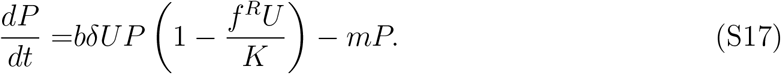

## 1.3 2-species bifurcation analysis

In the case of a single consumer feeding on a single resource, both the specialized (palatability) and the aggregate (recruitment) feedbacks simplify to the same dynamics characterized by a strong consumer Allee effect. In this case our model (eqns. 1–2) exhibits three equilibria: one with consumers extinct and resources at carrying capacity 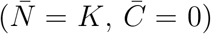 and two coexistence equilibria with 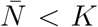 and 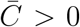. For simplicity, we assume *β* = 0. With both species present and *f*^*n*^ = *f*^*P*^ or *f*^*n*^ = *f*^*R*^, solving the equation of the consumer dynamics at 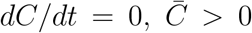 for the equilibrium resource abundance 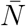 can give two positive coexistence equilibria

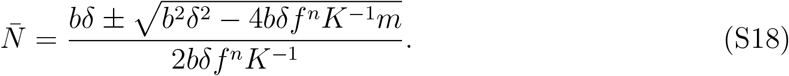

In our 2-dimensional system, the resource-only equilibrium has the Jacobian

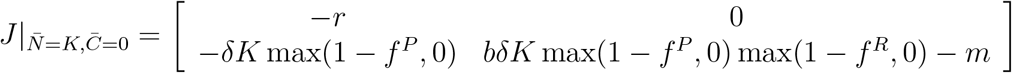

with eigenvalues *λ*_1_ = − *r* and *λ*_2_ = *bδK* max(1 – *f*^*P*^, 0) max(1 – *f*^*R*^, 0) – *m*. Giver *r* > 0, *λ*_1_ < 0, and the equilibrium is locally stable if *λ*_2_ < 0 and unstable otherwise. In both the palatability feedback model (*f*^*P*^ > 0, *f*^*R*^ = 0) and the recruitment feedback model (*f*^*P*^ = 0, *f*^*R*^ > 0), stability of the resource-only equilibrium then requires

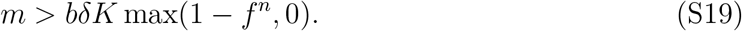

Existence of the coexistence equilibrium requires 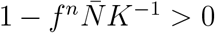, and so for simplicity we drop the max arguments. At the coexistence equilibrium the model-specific Jacobians are

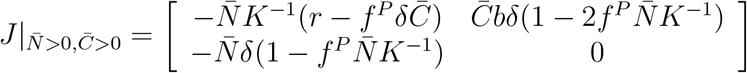

for specialized feedbacks and

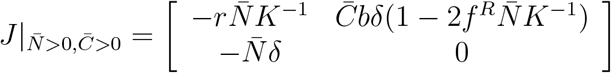

for aggregate feedbacks. Here we use the Routh-Hurwitz criteria (negative trace and positive determinant) for stability. In 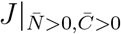 the bottom-left entries are always negative, such that for both feedbacks a positive determinant requires only 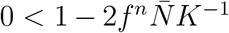. Substituting the lower 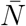 equilibrium from eqn. S18, stability of the coexistence equilibrium requires

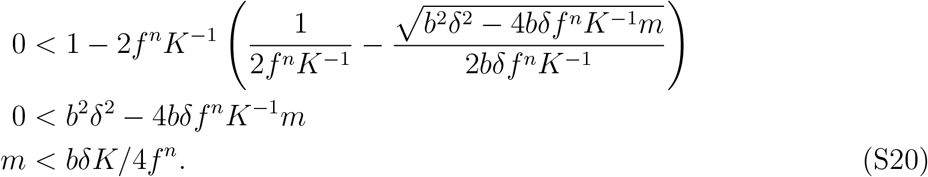

For palatability feedbacks, a negative trace in 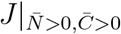 requires 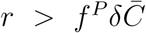. Solving *dN/dt* = 0 for 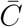 and substituting gives 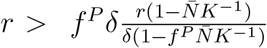, which simplifies to *f*^*P*^ < 1; in multi-species models interspecific competition among resources ensures 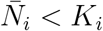, allowing coexistence equilibria with *f*^*P*^ > 1. Combining eqns. S19–S20, the range of mortality values leading to two stable states *D* is positive for *f* > 1*/*2 and given by

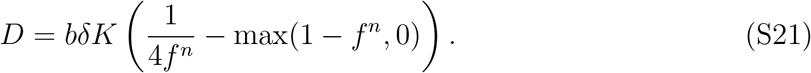

Critically, in the two-species case the range *D* of mortality levels leading to alternative stable states is identical for the palatability and the recruitment feedbacks because they share the same consumer dynamics. This analysis also highlights that consumer conversion efficiency (or the energetic value of resources), consumer grazing rate, and resource carrying capacity all increase the size of the bistability region. This reflects that stronger interactions promote bistability by reinforcing feedback loops. The change in hysteresis range with consumer grazing also underscores the importance of holding total consumption constant as we evaluate the effect of increased connectance on alternative stable states in our main analysis.

## Appendix B Dynamics with explicit stage structure

Here, we verify that the greater potential for aggregate feedbacks to produce alternative stable states remains in more complex models explicitly accounting for resource or consumer stage structure. We model food webs of 12 resource species, 8 consumer species, and either 12 resource juveniles (for specialized feedbacks) or 8 consumer juveniles (for aggregate feedbacks). For specialized feedbacks, we implement eqns. S7–S9 with mean parameter values *r* = 1, *δ* = 1.2, *b* = 1, *g* = 0.02, and *γ* = 0.5. For aggregate feedbacks, we implement eqns. S7–S9 with the simplifications *δ*_*J*_ = 0, *α*_*JC*_ = 0, and *α*_*JJ*_ = 0 which retain stage structure and the key cultivation-depensation feedback. We set the mean values of the remaining parameters to *r* = 1, *K* = 1.1, *δ*_*C*_ = 1.15, *β*_*C*_ = *b* = 1, *α*_*CJ*_ = *f* = 1.5, and *γ* = 0.3. Following the main text, for aggregate feedbacks we also incorporate the effects of each resource *i* on each consumer juvenile *j*, Ψ_*i,j*_, with *ψ* = 0.35. Finally for both models we incorporate density-dependent consumer mortality *β*_*P*_ = 0.075, among-resource competition *α* = 0.025, and trophic interactions Ω_*i,k*_ with *σ* = 0.5. With these parameters and a species heterogeneity of 0.4, we find qualitatively similar effects of connectance on the potential for alternative stable states in each model (Fig. S2).

## Appendix C Effects of specialization in recruitment and palatability interactions

Here, we expand our main analysis to consider how the potential for alternative stable states depends on both connectivity consumer diet as well as the degree of specialization in both classes of feedbacks. For recruitment feedbacks, greater degrees of specialization compared to those in our main analysis can arise when the survival of a consumer’s juveniles predominantly depends on the abundance of only a few resource species. With strong spatial sorting (or, equivalently, with relatively immobile adult consumers), specialization in recruitment feedbacks may also follow specialization in trophic interactions (i.e., Ψ_*i,k*_ = Ω_*i,k*_ *∀i, k*). For palatability feedbacks, our main analysis assumes that the palatability of one resource species depends only on that species’ density (i.e., high specialization). Here we relax this assumption by introducing juvenile resource *i* - adult resource *j* interactions Γ_*j,i*_ and, as with recruitment feedbacks, modify 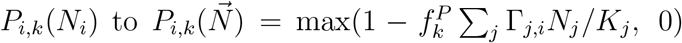. Palatability feedbacks involving size-dependent consumption may be less specialized when the proportion of vulnerable juveniles in a resource population depends on several resource species, as might occur when juvenile resources are outcompeted by the adults of multiple resource species rather than only by conspecific adults. Likewise, palatability feedbacks involving group defense may be less specialized when resource species form multi-species herds, schools, or plant stands to deter consumption.

Throughout, we define specialization *S* in interactions as the mean maximum weight of palatability dependence of one resource *i* on any other resource *j* (= 10^−1^ Σ_*i*_ max_*j*_(*Γ*_*j,i*_)) and, equivalently, the mean maximum weight of recruitment dependence of one consumer *k* on any resource *i* (= 10^−1^ Σ_*k*_ max_*i*_(*Ψ*_*i,k*_)). This metric is analogous to consumer electivity for its primary resource (Fig. 3). For both Ψ_*i,k*_ and Γ_*j,i*_, we draw interaction weights from a dirchlet distribution parameterized such that the largest entries in each column are, on average, *S*. For each model, we then measure (1) the hysteresis range of consumer mortality levels between the median points of consumer collapse and recovery and (2) state distinctiveness, measured as the maximum difference in consumer species richness between resource- and consumer-dominated food web states.

Our expanded analysis reiterates the greater distinctiveness and hysteresis range of alternative stable states with recruitment than with palatability feedbacks over all levels of connectance and feedback specialization (Fig. S3). In general, specialization tends to decrease the distinctiveness of alternative states for both feedbacks. We also find that high connectance and a very high low of specialization (*S <*0.2) can produce distinctive alternative stable states over a large range of mortality levels for palatability feedbacks. We consider this case uncommon, however, as juvenile resources nearing maturity are likely to have a stronger niche overlap with conspecific than with heterospecific adults. Finally, we find that recruitment feedbacks produce distinctive alternative stable states over a large range of mortality levels over all levels of connectance and feedback specialization, including scenarios where strong spatial structure where leads to Ψ_*i,k*_ = Ω_*i,k*_ ∀*i, k*.

## Appendix D Dynamics without consumer density dependence

Here we verify that the greater potential for aggregate feedbacks to produce alternative stable states remains without consumer density dependence, i.e., *β* = 0. To reduce among-consumer competition we model lower species heterogeneity of 0.1 (vs. 0.15 in the main text) and omit two consumer species (i.e., modeling 12 resources and 8 consumers). We find that the effects of connectance on the prevalence of alternative stable states for both feedbacks (Fig. S4) are analogous to the case with consumer density dependence (Fig. 3).

---

**Table S1:** Descriptions and mean values of parameters used in the base model (eqns. 1–2). Parameters without subscripts do not vary among species.

| Parameter | Description | Mean value |
| --- | --- | --- |
| $r_i$ | Resource $i$ growth rate | 1 |
| $K_i$ | Resource $i$ carrying capacity | 1.35 |
| $\alpha$ | Interspecific resource competition strength | 0.025 |
| $\delta_k$ | Consumer $k$ total grazing or consumption rate | 1.1 |
| $b_k$ | Conversion constant of resources to consumer $k$ recruits | 1 |
| $f_k^P$ | Decline in resource palatability with resource density for consumer $k$ | 1.2 |
| $f_k^R$ | Decline in consumer recruitment with resource density for consumer $k$ | 1.2 |
| $\sigma$ | Standard deviation of trophic interaction strengths in $\Omega_{i,k}$ | 0.5 |
| $\Omega_{i,k}$ | Proportion of total consumption a consumer $k$ allots to resource $i$ | varied |
| $\Psi_{i,k}$ | Strength of recruitment interaction between resource $i$ and consumer $k$ | |
| $\beta_k$ | Density-dependent consumer mortality | 0.15 |
| $m$ | Density-independent consumer mortality | varied |

**Figure S1:**
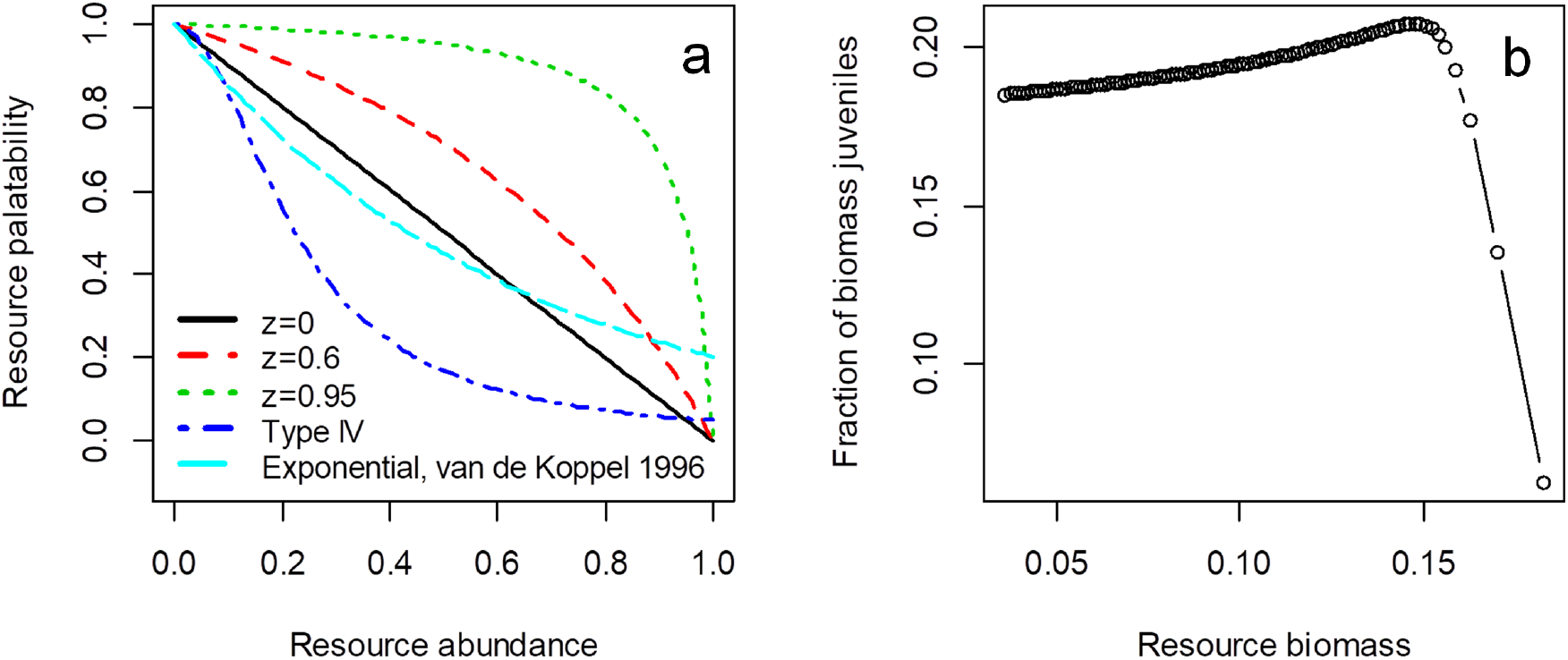
(a) Comparison of declines in resource palatability with abundance across levels of maturation sensitivity to competition (*z* in (1 – *A*)(1 – *zA*)^−1^), Type IV group defense functional response ((1+*ηN* ^2^)^−1^), and exponential declines in grazing as abundant vegetation impedes movement (exp(−*ηN*), *η* = 1.65 in van de Koppel et al. 1996). (b) Proportion of total equilibrium resource biomass represented by smaller (i.e., more palatable) juveniles across total equilibrium resource biomass in the model and parameterization of de Roos and Persson 2013 (Chapter 3, eqn. 24). Ranges of equilibrium biomass generated by increasing juvenile mortality to simulate different densities of a gape-limited consumer.

**Figure S2:**
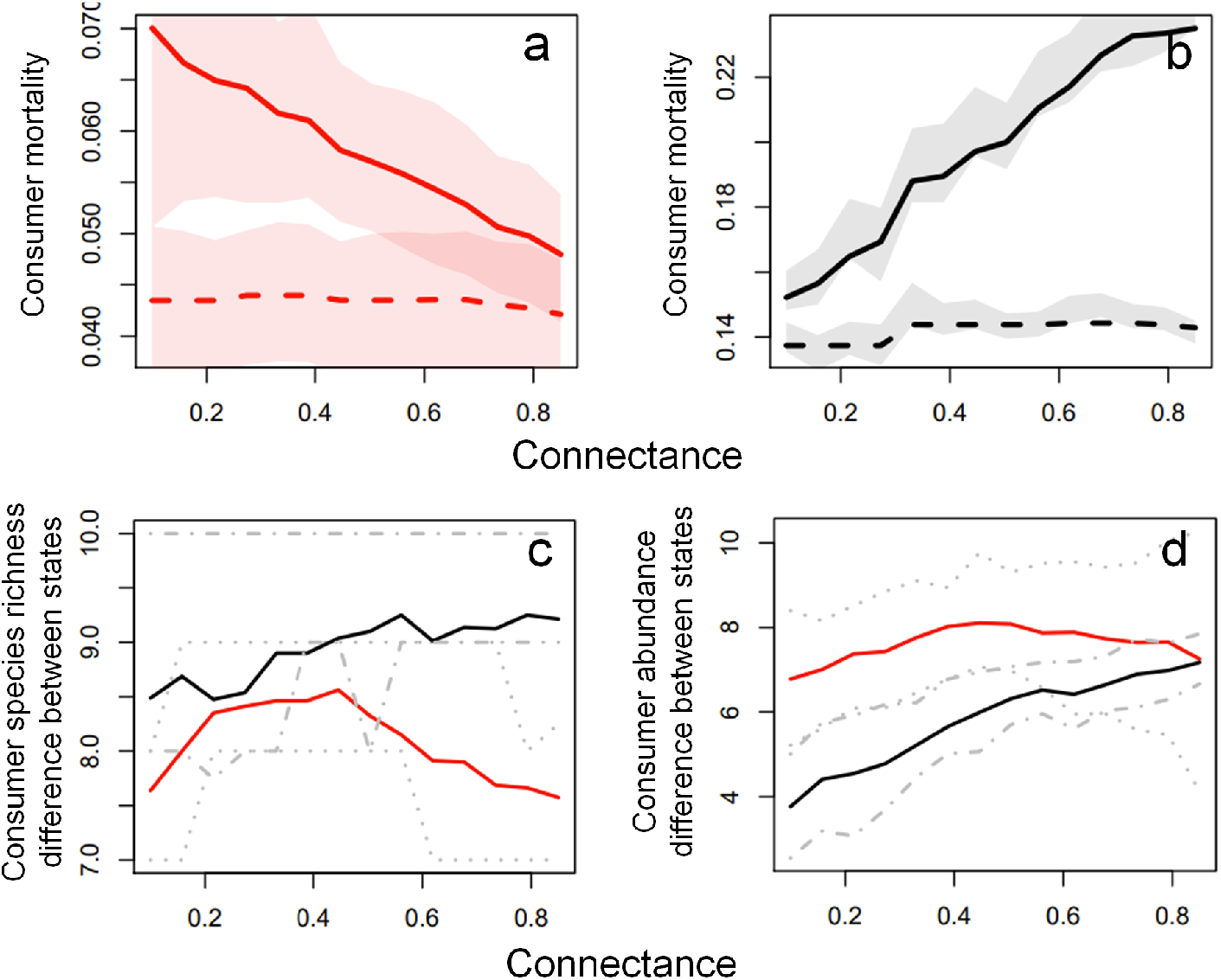
The potential for alternative stable states decreases with connectance for specialized feedbacks but increases with connectance for aggregate feedbacks in models with explicit stage structure. (a, b) Median thresholds at which consumer species collapse (solid lines) and recover (dashed lines) for specialized feedbacks (a, red colors) and aggregate feedbacks (b, black colors), with alternative stable states present between these thresholds. Shaded regions in (a, b) denote the interquartile range of each threshold. (c, d) Distinctiveness of consumer- and resource-dominated states, measured in species richness (c) and in total consumer abundance (d), with gray lines denoting interquartile ranges. All panels show results over 100 simulated food webs at each connectance level.

**Figure S3:**
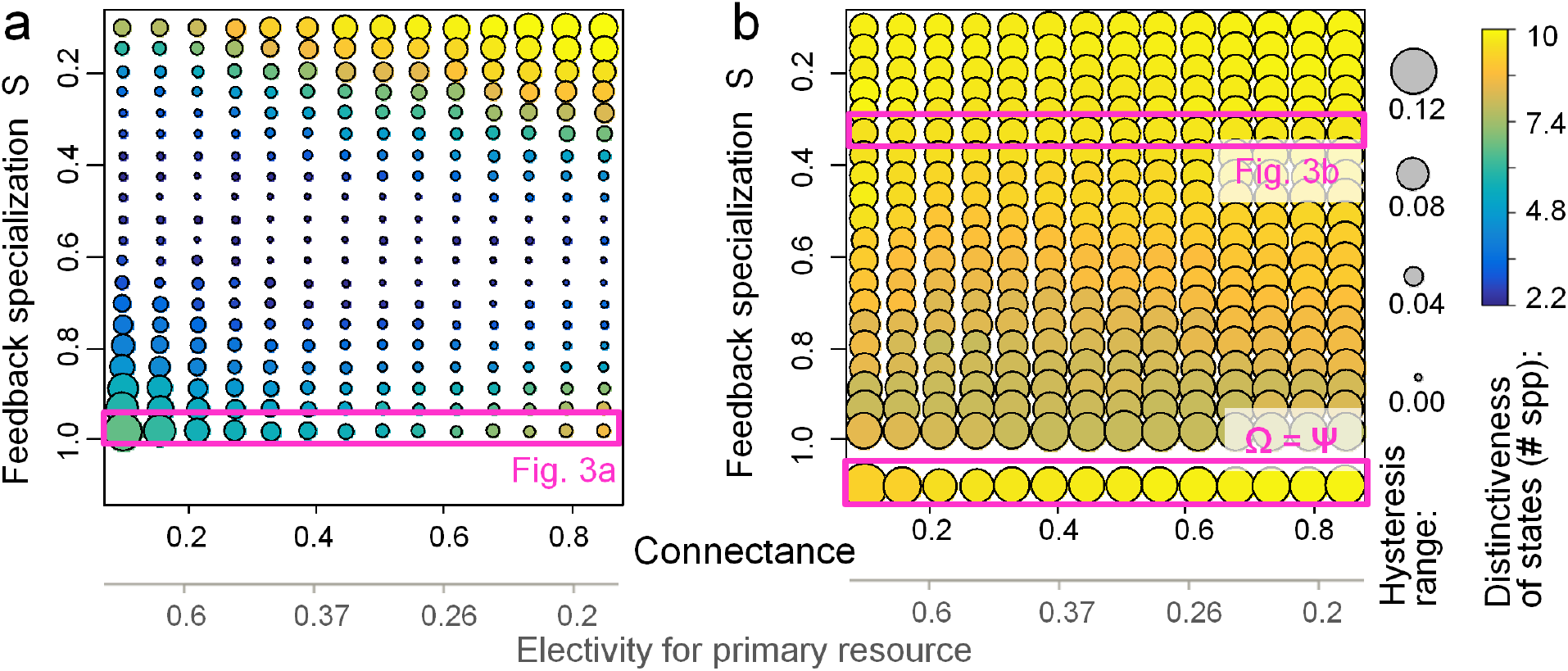
Palatability feedbacks (a) decrease the potential for alternative stable states compared with recruitment feedbacks (b) over all levels of connectance and feedback specialization *S*. Points at each connectance and specialization level denote the distinctiveness of alternative stable states (difference in consumer species richness between resource-and consumer-dominate states, point color) and the difference between the median mortality level of consumer collapse and the median mortality level of consumer recovery (point size), averaged over 120 simulated food webs. The secondary x-axis denotes the proportion of diet comprised by each consumer’s primary resource when resources are equally abundant (Ivlev electivity), and is analogous to feedback specialization.

**Figure S4:**
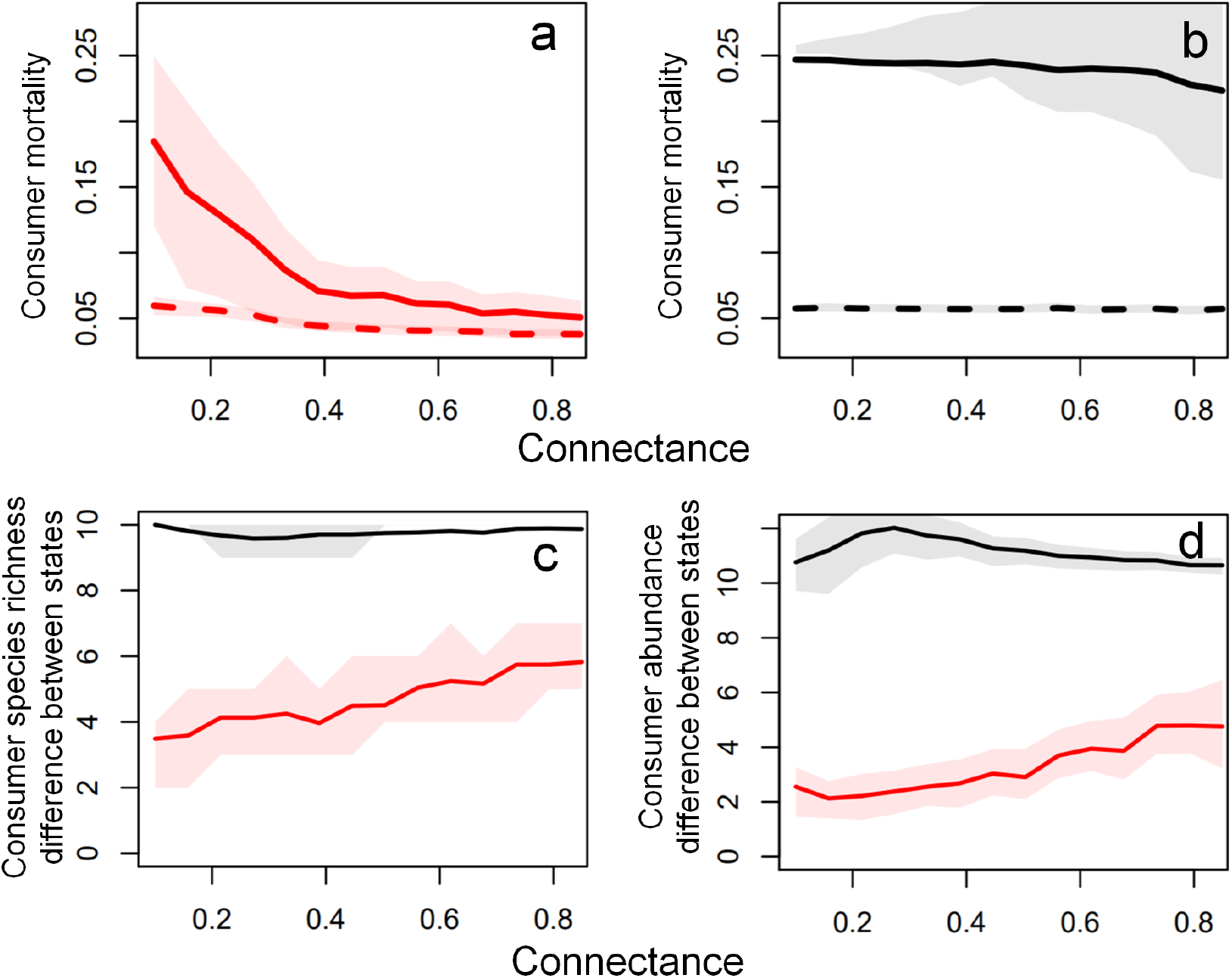
The potential for alternative stable states decreases with connectance for specialized feedbacks but increases with connectance for aggregate feedbacks in models without consumer density dependence. (a, b) Median thresholds at which consumer species collapse (solid lines) and recover (dashed lines) for specialized feedbacks (a, red colors) and aggregate feedbacks (b, black colors), with alternative stable states present between these thresholds. Shaded regions in (a, b) denote the interquartile range of each threshold. (c, d) Distinctiveness of consumer- and resource-dominated states, measured in species richness (c) and in total consumer abundance (d), with gray lines denoting interquartile ranges. All panels show results over 100 simulated food webs at each connectance level.

